# Palaeobiological inferences based on long bone epiphyseal and diaphyseal structure - the forelimb of xenarthrans (Mammalia)

**DOI:** 10.1101/318121

**Authors:** Eli Amson, John A Nyakatura

## Abstract

Trabecular architecture (i.e., the main orientation of the bone trabeculae, their number, mean thickness, spacing, etc.) has been shown experimentally to adapt with great accuracy and sensitivity to the loadings applied to the bone during life. However, the potential of trabecular parameters used as a proxy for the mechanical environment of an organism’s organ to help reconstruct the lifestyle of extinct taxa has only recently started to be exploited. Furthermore, these parameters are rarely combined to the long-used mid-diaphyseal parameters to inform such reconstructions. Here we investigate xenarthrans, for which functional and ecological reconstructions of extinct forms are particularly important in order to improve our macroevolutionary understanding of their main constitutive clades, i.e., the Tardigrada (sloths), Vermilingua (anteaters), and Cingulata (armadillos and extinct close relatives). The lifestyles of modern xenarthrans can be classified as fully terrestrial and highly fossorial (armadillos), arboreal (partly to fully) and hook-and-pull digging (anteaters), or suspensory (fully arboreal) and non-fossorial (sloths). The degree of arboreality and fossoriality of some extinct forms, “ground sloths” in particular, is highly debated. We used high-resolution computed tomography to compare the epiphyseal 3D architecture and mid-diaphyseal structure of the forelimb bones of extant and extinct xenarthrans. The comparative approach employed aims at inferring the most probable lifestyle of extinct taxa, using phylogenetically informed discriminant analyses. Several challenges preventing the attribution of one of the extant xenarthran lifestyles to the sampled extinct sloths were identified. Differing from that of the larger “ground sloths”, the bone structure of the small-sized *Hapalops* (Miocene of Argentina), however, was found as significantly more similar to that of extant sloths, even when accounting for the phylogenetic signal.

## INTRODUCTION

Bone structure is intensively studied in analyses concerned with functional anatomy because it is argued to be extremely plastic. While a genetic blueprint influences bone structure, it has been shown to adapt during life (and especially at an early ontogenetic stage) to its mechanical environment (Ruff et al. 2006). This was argued for trabecular bone, which reacts to loading with great accuracy and sensitivity (Barak et al. 2011). This was also argued for cortical bone, even though the latter is expected to be less plastic, at least in part due to its lower remodeling rate (see review of Kivell, 2016). Comparative studies focusing on either trabeculae or cortical structure intend to leverage this great plasticity to associate structural phenotypes to lifestyles or functional uses of a limb. This has been achieved in some analyses (as recently exemplified by Georgiou et al. 2018; Ryan et al. 2018; Tsegai et al. 2018) but not all of them (see review of Kivell 2016), suggesting that some confounding factors are likely to be at play, and more generally that the approach is limited. For trabecular bone in particular, important intraspecific variation has been documented (e.g., in *Pongo*; Tsegai et al. 2013; Georgiou et al. 2018). Nevertheless, the fact that some analyses successfully distinguished ecological groups might indicate that broad differences of bone structure among lifestyles can exceed, at least in some cases, individual variability. Because fossil bone cross-sections at mid-diaphysis have been produced for over a century and a half (Kolb et al. 2015), a large number of mid-diaphyseal data related to extinct taxa have been acquired, and successfully exploited for palaeobiological inferences (e.g., Germain & Laurin, 2005). Fossil three-dimensional (3D) trabecular architecture has been much less investigated, as, to our knowledge, only few studies have been published, which are all focussing on primates (DeSilva & Devlin 2012; Barak et al. 2013; Su et al. 2013; Skinner et al. 2015; Su & Carlson 2017; Ryan et al. 2018).

In general terms, it is assumed that the diaphysis of long bones tends to be exposed to mostly bending and torsion, and to a lesser extent axial compression (Carter & Beaupré 2001). On the other hand, the architecture of epiphyseal trabeculae is usually related to compressive and tensile strains (Biewener et al. 1996; Pontzer et al. 2006; Barak et al. 2011). Trabecular and cortical compartments are hence expected to have distinct mechanical properties, which do not necessarily co-vary. To combine them in a single analysis, it can therefore be argued that the structural parameters deriving from these two types of structures should be considered as distinct (univariate) variables. Because trabecular and cortical structures have independently yielded a functional signal, such a combined analysis could potentially help in our endeavours to associate a bone overall structure to a loading regime, and, eventually, a function. This combined analysis has previously been achieved, on extant taxa, via different approaches. Based on epiphyseal regions of interest (ROIs) and mid-diaphyseal sections, Shaw & Ryan (2012) examined both compartments in the humerus and femur of anthropoids (see also Lazenby et al. (2008) for handedness within humans). They measured individual trabecular and mid-diaphyseal parameters, but did not combine the latter in a single test. Another approach, termed ‘holistic analysis’ (Gross et al. 2014), was used in *Pan* and *Homo* whole bones or epiphyses, but parameters were not used conjointly to discriminate functional groups in the statistical assessment either. It is noteworthy, however, that Tsegai et al. (2017), also used this holistic analysis and performed a Principal Component Analysis (even though in that case the focus was on trabecular bone architecture and cortical bone thickness at the articular surface). Skinner et al. (2015) and Stephens et al. (2016) also used Gross et al. (2014)’s method, but focused on trabecular architecture only. This approach is particularly relevant for medium- to large-sized mammals such as *Pan* or *Homo*, for which the epiphyses include a complex trabecular architecture with distinct zones of different arrangement (such as the so-called vertical and horizontal trabecular columns in the femoral neck; Hammer 2010). One can note that an entirely different approach, not relying on the measurement of these parameters, but on micro-finite element analysis, was also applied to a primate (Huynh Nguyen et al. 2014). To our knowledge, epiphyseal trabecular and mid-diaphyseal parameters have never been combined in a functional analysis about non-primate taxa, and no analysis used both trabecular and cross-sectional parameters in the same discriminant test.

References to bone structure in “ground sloths”, *Megatherium* in particular, date back to the 19th century (Owen 1861). But it is only fairly recently that quantification of bone structure was performed (Straehl et al. 2013; see review of Amson & Nyakatura 2017). Straehl et al. (2013) examined compactness profile of a mid-diaphyseal section in the limb long bones of various extant and extinct xenarthrans. They found that most armadillos were characterized by a humeral mid-diaphysis that is relatively more compact than that of the femur. Subsequently, Amson et al. (2017a) studied the epiphyseal trabecular architecture in extant xenarthrans, and found that some parameters, the degree of anisotropy (DA) in particular, differed among functional categories.

Indeed, xenarthrans are marked by distinct lifestyles that can be used to define functional categories. Extant xenarthrans were categorized by Amson et al. (2017a) as fully arboreal and non-fossorial (extant sloths), intermediate in both fossoriality and arboreality (anteaters), or fully terrestrial and fossorial (armadillos), and several fossorial classes were recognized among the latter. Partly following their expectations, Amson et al. (2017a) recovered that the armadillos (and in particular the more highly fossorial ones) differ in their greater DA for instance. The latter can be expected to be associated with the presence of one main loading direction in these highly fossorial taxa (as opposed to various equally marked directions in taxa for which the forelimb is arguably facing a less stereotypical main loading). Similarly, one could expect those taxa of which the long bone in question experiences one main bending direction to be characterized by a more elliptical cross-sectional shape at mid-diaphysis (CSS, see below), with the section’s major axis aligned along that direction (as the major axis indicates the direction of the greatest bending rigidity; Ruff & Hayes 1983). Because no significant differences were recovered in the mid-diaphyseal global compactness between fossorial and non-fossorial talpid moles (Meier et al. 2013), it seems that a simple relation between this parameter and a loading scheme associated with fossorial activity should not be expected (see also Straehl et al. 2013).

For extant xenarthrans, the functional categories mentioned above mostly match the phylogeny, i.e., most categories are aggregated into clades. However, this is likely not true if one includes the extinct xenarthrans, the “ground sloths” in particular, because their lifestyle was interpreted as different from that of their closest relatives, the “tree sloths”. Lifestyle reconstruction of extinct xenarthrans dates back to the 18th century (see review of Amson & Nyakatura 2017). Various methods were employed to infer the lifestyle of extinct xenarthrans. So far, they all relied on bone (and tooth) gross morphology, involving approaches such as comparative functional morphology (Amson et al. 2015a), biomechanical modelling (Fariña & Blanco 1996) or muscle reconstruction (Toledo et al. 2013). This was found to be challenging, partly because of the lack of modern analogues for some taxa (Vizcaíno et al. 2017), and partly because of the autapomorphic nature of several of the xenarthran traits. This, along with the fact that functional categories mostly match phylogeny, makes disentangling the phylogenetic and functional signals difficult (Amson et al. 2017a). Bone structure was argued to be extremely plastic and found in xenarthrans in particular to be mostly devoid of phylogenetic signal (and when a significant signal is found, it is likely due to the matching between functional categories and clades; Amson et al. 2017a). The ecophenotypic nature of bone structure traits (which are defined as “biomechanically informative phenotypically plastic”; Ryan et al. 2018) is the rationale behind the present endeavour.

The aim of this study is to quantify bone diaphyseal and trabecular structure in “ground sloths” in order to infer their lifestyle. Given the disparate gross morphology of xenarthrans (e.g., the humerus is extremely slender in extant sloths and particularly stout in most armadillos, see Mielke et al. 2018a), we believe that studying easily comparable and arguably ecophenotypic traits such as bone structure parameters is highly relevant for this purpose. Extant sloths represent but a remnant of the overall diversity of Tardigrada (also termed Folivora or Phyllophaga; Delsuc et al. 2001), and the two extant genera, *Bradypus* (three-toed sloths) and *Choloepus* (two-toed sloths), most likely acquired their highly derived lifestyle convergently (Nyakatura 2012; Coutier et al. 2017). Most aspects of the biology of “ground sloths” exceed in disparity those of their extant kin. They were found from Alaska (Stock 1942) to southernmost South America (and potentially Antarctica; Carlini et al. 1990; Gelfo et al. 2015) Various feeding habits, such as bulk-feeding or selective-feeding are purported (Bargo & Vizcaíno 2008). The lifestyle of most extinct sloths is reconstructed as terrestrial (but see *Thalassocnus* for an (semi-)aquatic lineage; Amson et al. 2015b). Furthermore, some “ground sloths” contrast with their extant relatives in reaching large body sizes (up to several tones; Fariña et al. 1998).

The fossil record of early (Palaeocene-Eocene) xenarthrans and especially that of sloths, is rather poor (Gaudin & Croft 2015). It is therefore hard to reconstruct the ancestral lifestyle of Tardigrada, and more generally Xenarthra. To date, no extinct sloths have been reconstructed to have had a suspensory posture and locomotion resembling their extant kin (Pujos et al. 2012). But, because their gross anatomy was considered as similar to that of extant anteaters, Matthew (1912) argued that *Hapalops*, for instance, was partly arboreal. Such a lifestyle was of course not considered for larger taxa (but see translation of Lund in Owen (1839) for an early opposite view). However, digging capabilities, as well as bipedal stance and/or locomotion, was proposed for several medium-sized (e.g., *Glossotherium*) to giant-sized (e.g., *Megatherium*) “ground sloths” (Bargo et al. 2000; Patiño & Fariña 2017). For the present analysis, we were able to sample small-sized as well as large-sized “ground sloths.” The estimated body sizes of the latter exceed that of extant xenarthrans by two orders of magnitude (see below for body mass estimates). Because this has already been pointed out as a challenge for the reconstruction of extinct xenarthrans’ lifestyles (Vizcaíno et al. 2017), and because size might be correlated to at least some bone structure parameters, potential challenges inherent to the taxa and parameters we studied will be discussed.

## MATERIAL AND METHODS

### Specimen and scanning procedure

The dataset of Amson et al. (2017a), which consists of extant skeletally mature wild-caught xenarthrans, was extended by several extinct sloths roughly spanning the whole body size range of the group: the small-sized (ca. 38 kg; Bargo et al. 2012) *Hapalops* sp. (Santa Cruz Formation, Early Miocene, ca. 17 Ma; Perkins et al. 2012), the medium-sized (ca. 200 kg; Smith et al. 2003) *Valgipes bucklandi* (Lagoa Santa, Brazil, Pleistocene; the sampled specimen MNHN.F.BRD29 is labelled *Ocnopus gracilis*, which is now viewed as a junior synonym; Cartelle et al. 2009), *Scelidotherium leptocephalum* (ca. 1000 kg; Vizcaíno et al. 2006) and *Glossotherium robustum* [ca. 1200 kg (Vizcaíno et al. 2006); both from ‘Pampean’, Argentina and Tarija, Bolivia, both Pleistocene], as well as the large-sized *Lestodon armatus* [ca. 3200 kg (Vizcaíno et al. 2006); ‘Pampean’, Argentina, Pleistocene] and *Megatherium americanum* [ca. 4000 kg (Fariña et al. 1998); ‘Pampean’, Argentina, Pleistocene]. The sampled specimens are skeletally mature (a few specimens showed a remnant of epiphyseal line, see below) and did not present apparent bone diseases (which were also criteria of selection for the extant species, see Amson et al. 2017a). All fossils were scanned (micro computed tomography, μCT) using a vltomelx 240 L system (GE Sensing & Inspection Technologies Phoenix Xlray) at the AST-RX platform of the Museum national d’Histoire naturelle (Paris, France). According to the methodology and results of Amson et al. (2017a), we focused our data acquisition of the trabecular parameters on the humeral head and radial trochlea regions of interest (ROIs; see below). Mid-diaphyseal parameters were acquired for these two bones and for the third metacarpal (Mc III) in all species, when available. See Table 1 for the list of skeletal elements sampled for each extinct species, along with ROIs for which data were successfully acquired [see also Amson et al. (2017a), for sample size and scanning procedure of the extant species specimens]. For the included specimens, scanning resolution ranged from 0.03 to 0.123 mm (depending on the size of the specimens). Relative resolution, used to assess if the employed resolution is adequate to analyse trabecular bone (mean trabecular thickness divided by resolution) ranged from 5.1 to 11.5 pixels/trabecula. This is considered as appropriate (Sode et al. 2008; Kivell et al. 2011; Mielke et al. 2018b). Scanning resolution (and relative resolution for the trabecular ROIs) for each specimen can be found in Supplementary Online Material (SOM) 1. For this first endeavour of palaeobiological reconstruction of “ground sloths” lifestyle based on bone diaphyseal and trabecular structure, we compared the parameters yielded by the fossils to those of the extant specimens, using the same lifestyle categories as defined by Amson et al. (2017a), i.e., the fully arboreal extant sloths, intermediate anteaters, and fully terrestrial and fossorial armadillos.

**Table 1.** List of fossils with type of data acquired for each bone.

| Species | Specimen number | Data type |  |  |
| --- | --- | --- | --- | --- |
|  |  | Humerus | Radius | Mc III |
| *Hapalops* sp. | MNHN.F.SCZ166 | - | 72%MD; 100%TA | - |
| *Hapalops* sp. | MNHN.F.SCZ164 | 35%MD; 72%TA; (39%TA) | - | - |
| *Hapalops* sp. | MNHN.F.SCZ162 | 50%MD; 35%MD; (39%TA) | - | - |
| *Lestodon armatus* | MNHN.F.PAM754 | - | 50%MD |  |
| *Lestodon armatus* | MNHN.F.PAM755 | - | - | 50%MD |
| *Lestodon armatus* | MNHN.F.PAM95 | 100%TA | - | - |
| *Glossotherium robustum* | MNHN.F.PAM756 | QO | 50%MD; 100%TA | - |
| *Glossotherium robustum* | MNHN.F.PAM141 | - | - | 50%MD |
| *Scelidotherium leptocephalum* | MNHN.F.PAM236 | 50%MD | - | - |
| *Valgipes bucklandi* | MNHN.F.BRD29 | - | - | 50%MD |
| *Megatherium americanum* | MNHN.F.PAM753 | - | - | 50%MD |
| *Megatherium americanum* | MNHN.F.PAM758 | - | QO | - |

Footnotes. Abbreviations: 'n'-MD, mid-diaphyseal data, with 'n' the position of the sampled cross-section expressed as the length percentage from the proximal end; 'n'-TA, trabecular architecture data, with 'n' the cropping coefficient that was used, if any (see Material and Methods section); QO, only qualitative observations were performed.

### Qualitative observation of the diaphyseal structure

Raw image stacks were visualized with the Fiji package (ImageJ2 v. 1.51n and plugins; Schindelin et al. 2012, 2015; Schneider et al. 2012). The ‘Orthogonal Views’ routine was used to compute longitudinal sections. Sedimentary matrix prevented satisfying segmentation for some specimens but at least some qualitative observations were possible for all specimens (see Table 1).

### Trabecular parameters

We followed the methodology of Amson et al. (2017a), which involves the use of the BoneJ plugin (Doube et al. 2010) for Fiji. In brief, bones were first placed in the same standard orientation. Then, ROIs were selected in the centre of the studied epiphyses, with the ‘Fit Sphere’ routine of BoneJ (see Amson et al. 2017a: fig. 2 and Additional files 3, 4). ROI were selected to be as large as possible but without including cortical bone. We used the ‘Orthogonal Views’ routine of Fiji to ascertain that the centre of the ROI was precisely located at the centre to the studied epiphysis along the mediolateral, anteroposterior, and proximodistal directions. The resulting substack was then thresholded (‘Optimise Threshold> Threshold Only’ routine) and purified (‘Purify’ routine). Finally, trabecular parameters were measured. Given the results of Amson et al. (2017a), we focused on the degree of anisotropy (DA), main direction of the trabeculae (MDT), bone volume fraction, BV/TV, connectivity density (Conn.D), trabecular mean thickness (Tb.Th), trabecular mean spacing (Tb.Sp), bone surface area (BS). Other trabecular parameters routinely acquired, however, can also be found in SOM 1.

For some specimens, the lack of contrast between bone and the sedimentary matrix prevented accurate bone segmentation (see Table 1). Thresholding (see above) was successfully performed for the rest of the specimens; some of the latter, however, required manual removal of a few sedimentary particles (using the un-thresholded stack to recognize them).

The humerus of two specimens of *Hapalops* showed a slight remnant of epiphyseal line. A smaller ROI was hence defined to exclude this line (which would have biased the measurements) by cropping isometrically (in 3D) the substack (custom ImageJ script, SOM 2). The cropping coefficient (MNHN.F.SCZ162: 39%; MNHN.F.SCZ164: 72%) was then applied to the whole dataset and trabecular parameters were acquired anew. The means of the latter were compared to the initial parameters. For the dataset cropped at 72%, differences were found as minor (similar MDT; ΔDA = 3%; ΔBV/TV < 1%; ΔConnD <1%), while for the dataset cropped at 39%, differences were more important (MDT of opposing direction; ΔDA, 13%; ΔBV/TV = 2%; ΔConnD, 4%). Because it was exceeding a difference of 5% for at least one parameter value, we did not analyse further the latter dataset (and excluded MNHN.F.SCZ162 from the analysis of trabecular architecture). A remnant of epiphyseal line was also observed in *Glossotherium robustum* MNHN.F.PAM756, but in its case only qualitative observations were made.

### Mid-diaphyseal parameters

The same standardly oriented μCT-scan stacks (see above) were used for the acquisition of mid-diaphyseal parameters. Using Fiji, a cross-section was selected at mid-diaphysis; the latter was defined as the midpoint between most proximal and most distal points of either articular surfaces. Several sampled fossils did not preserve the mid-diaphysis. To compare them to the rest of the specimens, the latter were re-sampled at the level closest to mid-diaphysis preserved by each of those fossils (as assessed by superimposition with a complete specimen of the same species; MNHN.F.CSZ164 (humerus): 35% from proximal end; MNHN.F.CSZ166 (radius): 72% from the proximal end; see Table 1).

Once the diaphyseal cross-sections were selected, they were thresholded automatically (see above), but we manually checked the resulting image, which, in a few instances, required a manual correction of the levels. The whole sectional area (WArea), global compactness (GC; both acquired with a custom ImageJ script, SOM 3), and cross-sectional parameters of the ‘Slice Geometry’ routine of BoneJ (Doube et al. 2010) were acquired. For the following analyses, we focused on cross-sectional area (CSA) and the ratio of second moment of area around major to minor axes (Imax/Imin), also termed cross-sectional shape (CSS). If the ratio is close to one, CSS will usually be roughly circular. Values above one will entail increasingly elliptical shapes. The other diaphyseal parameters, however, can also be found in SOM 1. Because, if normalized with WArea (see below), CSA would be redundant with GC, it will only be used as a potential body size proxy.

### Statistics

The statistical analysis was performed using R version 3.4.3. Amson et al. (2017a) accounted for size effects by computing a phylogenetically informed linear regression for each parameter, against a size proxy. If the regression was found as significant, its residuals were used as the ‘size-corrected’ parameter. But the size of “ground sloths”, well exceeding for most of them that of extant xenarthrans, could bias such a procedure. Indeed, the slightest error on the regression coefficients estimation would likely involve drastically different residuals for those outlying taxa (see also Discussion). We therefore favoured, for the present analysis, to normalize those parameters that have a dimension by dividing the trait value by a body size proxy (raised to the same dimension). As body size proxies, we considered the specimen-specific TV (for trabecular parameters) and WArea (mid-diaphyseal parameters) or body mass (BM; species averages, because unknown for most collection specimens). Species body masses were taken from the AnAge database (Tacutu et al. 2013) and additional sources when necessary (Vizcaíno et al. 1999; Hayssen 2010; Abba & Superina 2016; Smith & Owen 2017) for the extant species and from the specific sources mentioned above for the extinct taxa. The coefficient of determination of regressions against a parameter well known to correlate with size (Tb.Th for trabecular parameters and CSA for mid-diaphyseal parameters) indicated that BM was more representative of the sample variance for the trabecular parameters, while it was WArea in the case of mid-diaphyseal parameters. They were accordingly used as body size proxies in the subsequent analyses.

Besides univariate comparisons, we performed linear discriminant analyses to infer the most likely lifestyle of extinct species. Both trabecular and mid-diaphyseal parameters of the humerus and radius were conjointly used in these analyses (parameters from the Mc III were not included because of their lack of discrimination power, see Results). To account for the great body size disparity of the studied taxa, it is the ‘size-normalized’ parameters that were used (raw value divided by the relevant body size proxy if parameter not dimensionless, see above). One analysis per extinct taxon was performed, because we were not able to acquire all parameters for each of them (depending on the successfully processed skeletal elements and ROIs, see Table 1). To phylogenetically inform these analyses, we used the function pFDA (Motani & Schmitz 2011; latest version available on github.com/lschmitz/phylo.fda). This ‘phylogenetic flexible discriminant analysis’ uses the optimised value of Pagel’s Lambda to account for the phylogenetic signal (Pagel 1999). As implemented here, the latter can span from 0 to 1, respectively denoting absence of phylogenetic signal and trait evolution consistent with a Brownian motion model of evolution. The rest of the pFDA works as a ‘traditional’ discriminant analysis. The training data, stemming in our case from the extant xenarthrans, were classified according to the three main lifestyles, i.e., ‘armadillo’, ‘anteater’, and ‘extant sloth’. The test data relates to the sampled extinct sloths. If not already normally distributed (as indicated by a Shapiro test), the parameters were log-transformed (and Shapiro tests were run again to confirm normality). Collinear variables (highly correlated variables as indicated by a correlation above 0.9) were excluded.

The timetree used to phylogenetically inform the tests was based on that used by Amson et al (2017a) (which is based on Gibb et al. 2016), and was completed with the extinct taxa. The relationships between the main clades follow Amson et al. (2017b). The split between Mylodontidae (represented by *Lestodon*) and the other Eutardigrada (all sloths but *Bradypus*) was set according to the age of the oldest fossil pertaining to the clade (*Octodontotherium*, ca. 29 Ma; Flynn & Swisher 1995; Kay et al., 1998) and is thus conservative (Fig. 1). But one can note that this age is roughly as old or older that the recent molecular estimations of the divergence time between the two genera of extant sloths (Slater et al. 2016; Delsuc et al. 2018). The age of divergence between *Lestodon* and *Glossotherium* was set according to the age of *Thinobadistes* (Hemphillian, ca. 9 Ma; Woodburne 2010), which is more closely related to *Lestodon* than *Glossotherium* according to Gaudin (2004). Extinct sloths were placed according to their known geological ages (see above; for Pleistocene taxa, a relatively young age of 0.1 Ma was arbitrarily given. Length of the branches leading to nodes of unknown ages, which are in direct relation to extinct taxa, and from these to terminal extinct taxa, were arbitrarily set to 1 and 0.1 Ma, respectively. Caution should be taken regarding the phylogenetic scheme used herein, because recent developments (yet to be published) in phylogenetic analyses of xenarthrans, which involve ancient DNA, might imply significant alterations of our understanding of sloths’ systematics (R.D.E. MacPhee, pers. comm., 2018).

**Figure 1.**
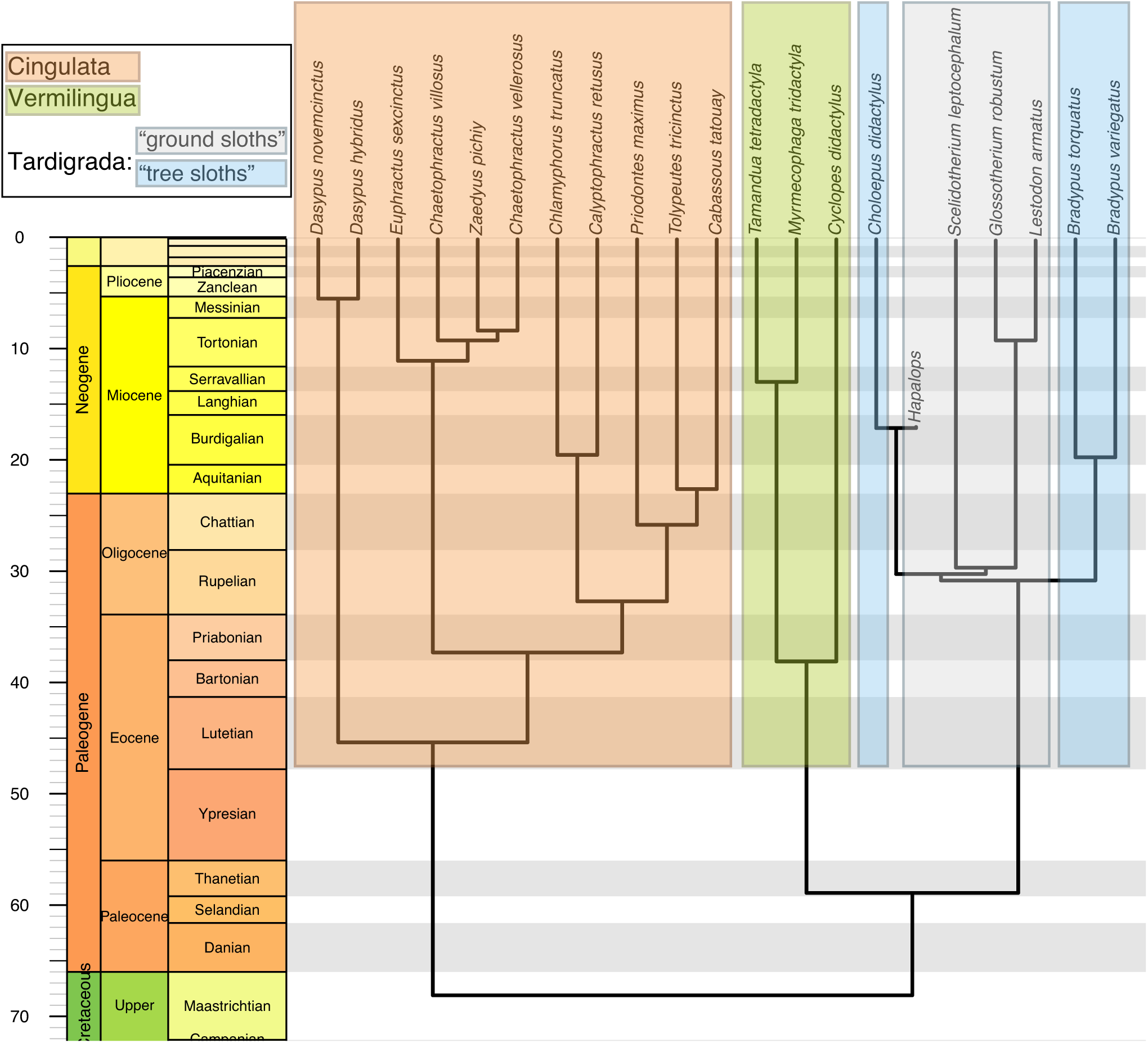
Timetree depicting the time-calibrated phylogenetic relationships of the xenarthrans included in the phylogenetically flexible linear discriminant analyses. See Material and Methods section for the sources used to build the timetree.

## RESULTS

### Qualitative observations of diaphyseal structure

In the humerus of small armadillos and anteaters, the medullary cavity is mostly devoid of spongy bone (with just a few isolated trabeculae, e.g., *Chaetophractus vellerosus* ZSM-1926-24, Fig. 2A; *Cyclopes didactylus*, ZMB_MAM_3913). In larger members of these clades, the medullary cavity is filled throughout the proximodistal length of the diaphysis by a more or less dense spongiosa (e.g., *Priodontes maximus* ZSM-1931-293; *Myrmecophaga tridactyla*, ZMB_MAM_102642; Fig. 2B-C). In extant sloths, a spongiosa can be observed in most of the diaphysis (*Bradypus*; n=4) or throughout its length (*Choloepus*, Fig. 2E; n=4), but a central region free of trabeculae subsists. The medullary cavity of the whole diaphysis is full of spongy bone in *Glossotherium* (n=1; Fig. 2F). It is nearly full in *Scelidotherium*, with just a small central free region subsisting (n=1). For *Hapalops*, a clear assessment cannot be given due to the preservation of the specimens at hand (MNHN.F.SCZ162 seems to show a free medullary cavity, but MNHN.F.SCZ164, which only preserves the proximal third of bone, shows a medullary cavity full of spongy bone). The whole diaphysis of the larger sloths *Megatherium* and *Lestodon* were not observed, but it is noteworthy that their epiphyses are filled with dense spongiosa (each n=1).

**Figure 2.**
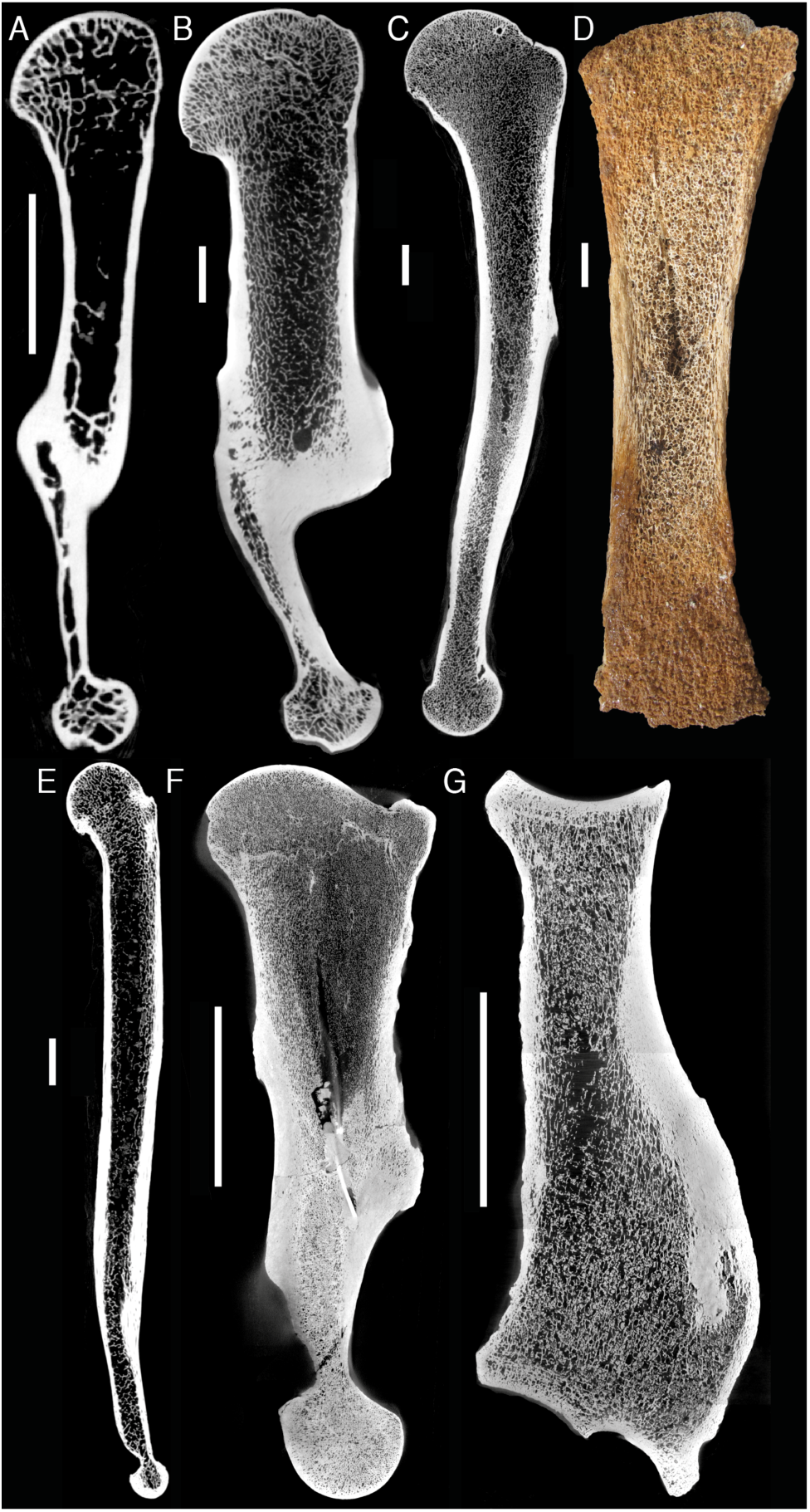
Qualitative observations of diaphyseal structure in xenarthrans. Longitudinal sections of humeri (A-C, E-F, all from CT-scans), tibia (D, ‘natural’ section), and radius (G, from CT-scan). A, *Chaetophractus vellerosus* (ZSM 1926-24); B, *Priodontes maximus* (ZSM 1931-293); C, *Myrmecophaga tridactyla* (ZMB_MAM_77025); D, *Nothrotherium maquinense* (MCL 2821); E, *Choloepus didactylus* (ZMB_MAM_35825); F, *Glossotherium robustum* (MNHN.F.TAR 767); G, *Lestodon armatus* (MNHN.F.PAM 754). Scale bars: A-E, 1 cm; F-G, 10 cm.

The radius of extant xenarthrans shows the same pattern as the humerus. In *Glossotherium*, *Lestodon*, and *Megatherium*, the medullary cavity of the whole radial diaphysis is essentially full of spongy bone (Fig. 2G; no data for *Hapalops* for which the entire radial epiphysis could not have been sampled).

### Univariate comparisons

The structure of the Mc III of extant species did not differ notably among the lifestyle categories (Fig. 3AB; Table 2). There is only a tendency for the anteaters and armadillos to have a more compact mid-diaphysis (Fig. 3A). Mc III structure was therefore not further studied, and not included in the discriminant analyses (see below). One can note, however, that some armadillos have an outlyingly high CSS (i.e., very elliptic cross-section) at mid-diaphysis (Fig. 3B; the single most elliptic value is found in the subterranean *Calyptophractus retusus* ZSM-1961-316). A great disparity of CSS at this location is found in extinct sloths, with the value of *Valgipes* falling among the outlying armadillos just mentioned, and that of *Megatherium* being the single lowest (i.e., most circular cross-section).

**Table 2.**
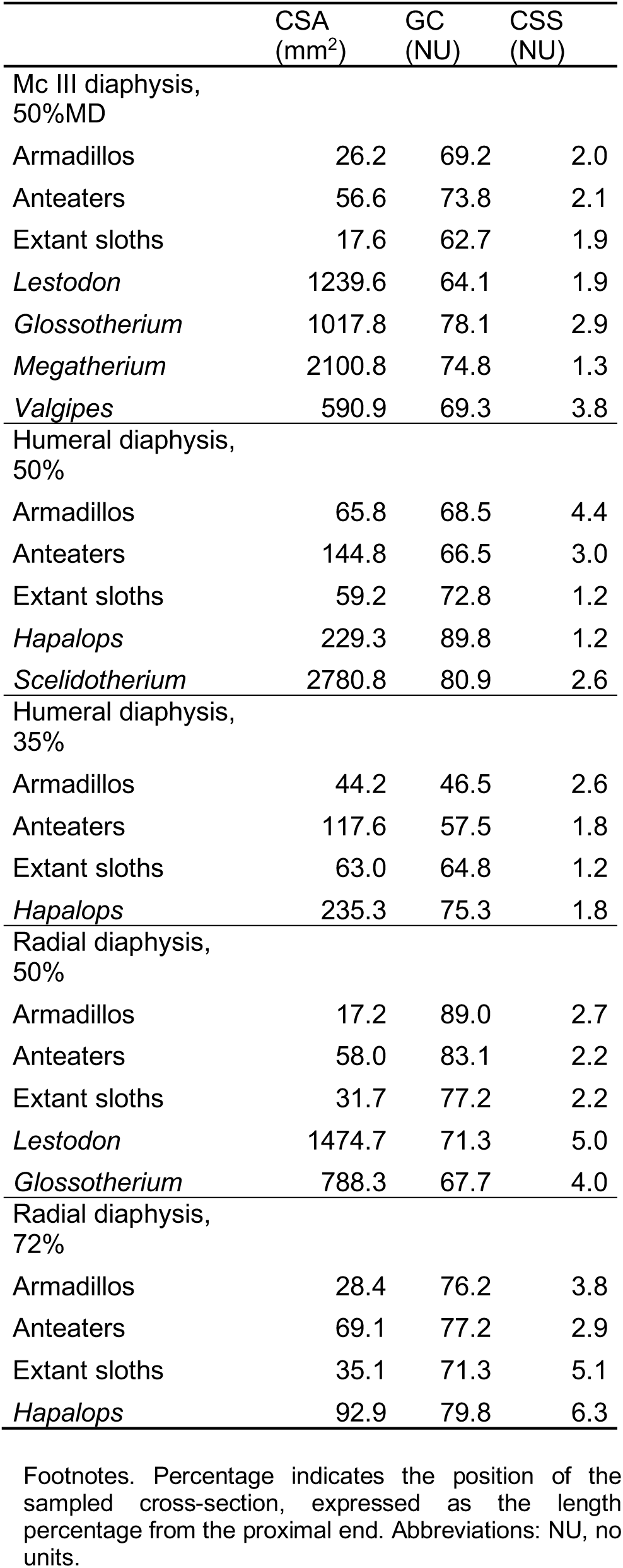
Mean values of diaphyseal parameters of interest for each lifestyle category and extinct taxon.

|  | CSA<br>(mm<sup>2</sup>) | GC<br>(NU) | CSS<br>(NU) |
| --- | --- | --- | --- |
| Mc III diaphysis,<br>50%MD |  |  |  |
| Armadillos | 26.2 | 69.2 | 2.0 |
| Anteaters | 56.6 | 73.8 | 2.1 |
| Extant sloths | 17.6 | 62.7 | 1.9 |
| *Lestodon* | 1239.6 | 64.1 | 1.9 |
| *Glossotherium* | 1017.8 | 78.1 | 2.9 |
| *Megatherium* | 2100.8 | 74.8 | 1.3 |
| *Valgipes* | 590.9 | 69.3 | 3.8 |
| Humeral diaphysis,<br>50% |  |  |  |
| Armadillos | 65.8 | 68.5 | 4.4 |
| Anteaters | 144.8 | 66.5 | 3.0 |
| Extant sloths | 59.2 | 72.8 | 1.2 |
| *Hapalops* | 229.3 | 89.8 | 1.2 |
| *Scelidotherium* | 2780.8 | 80.9 | 2.6 |
| Humeral diaphysis,<br>35% |  |  |  |
| Armadillos | 44.2 | 46.5 | 2.6 |
| Anteaters | 117.6 | 57.5 | 1.8 |
| Extant sloths | 63.0 | 64.8 | 1.2 |
| *Hapalops* | 235.3 | 75.3 | 1.8 |
| Radial diaphysis,<br>50% |  |  |  |
| Armadillos | 17.2 | 89.0 | 2.7 |
| Anteaters | 58.0 | 83.1 | 2.2 |
| Extant sloths | 31.7 | 77.2 | 2.2 |
| *Lestodon* | 1474.7 | 71.3 | 5.0 |
| *Glossotherium* | 788.3 | 67.7 | 4.0 |
| Radial diaphysis,<br>72% |  |  |  |
| Armadillos | 28.4 | 76.2 | 3.8 |
| Anteaters | 69.1 | 77.2 | 2.9 |
| Extant sloths | 35.1 | 71.3 | 5.1 |
| *Hapalops* | 92.9 | 79.8 | 6.3 |

Footnotes. Percentage indicates the position of the sampled cross-section, expressed as the length percentage from the proximal end. Abbreviations: NU, no units.

**Figure 3.**
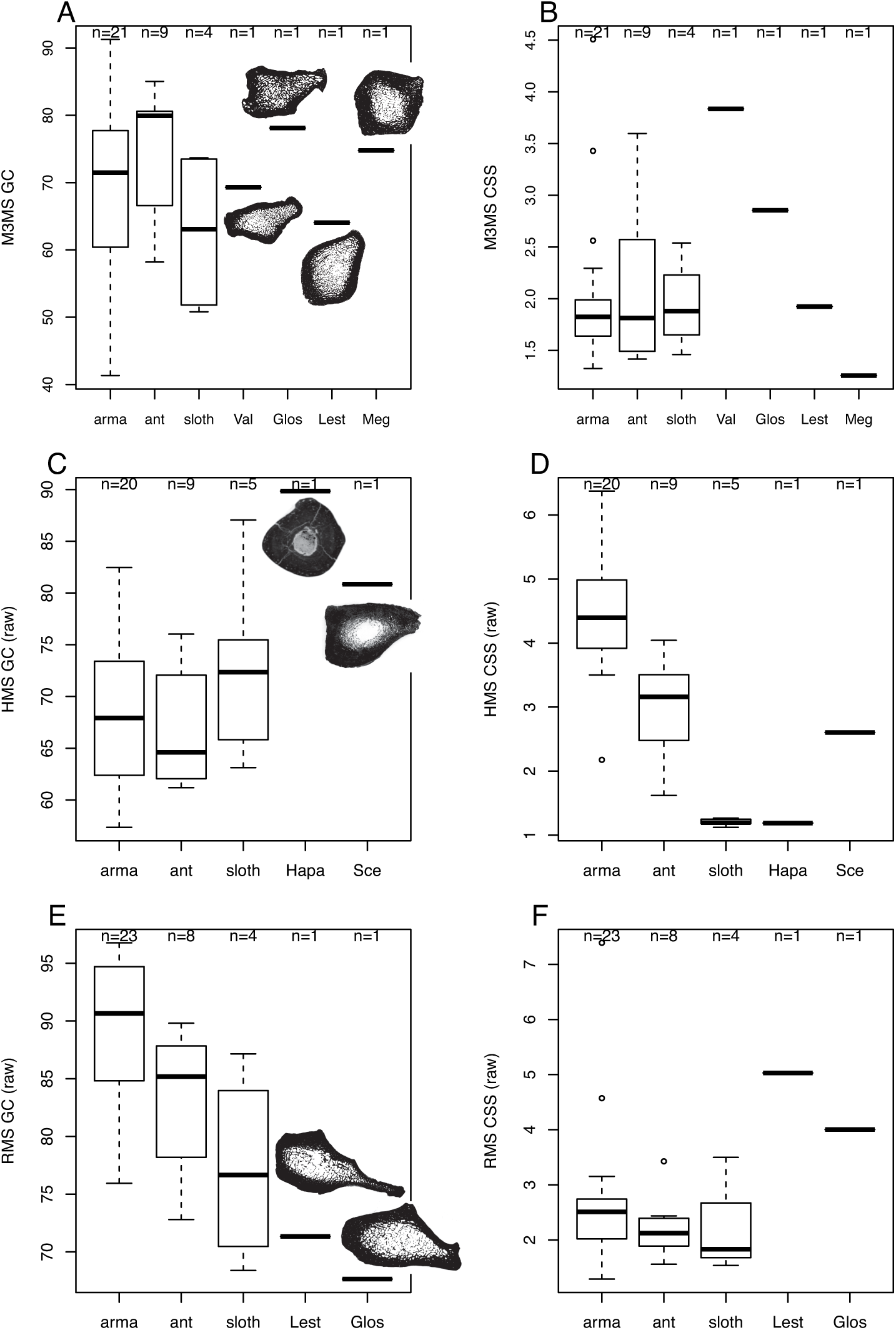
Univariate comparisons of middiaphyseal parameters. A, Mc III Global Compactness (GC); B, Mc III cross-sectional shape (CSS); C, humeral GC; D, humeral CSS; E, radial GC; F, radial CSS. Thresholded middiaphyseal virtual sections are depicted for the extinct sloths. Abbreviations: ant, anteaters; arma, armadillos; Glos, *Glossotherium*; Hapa, *Hapalops*; Lest, *Lestodon*; Meg, *Megatherium*; Sce, *Scelidotherium*; sloth, extant sloths.

The humeral diaphysis in *Hapalops* is remarkably compact. At mid-diaphysis (n=1), it features the highest GC value of the whole dataset (Fig. 3C; Table 2). At 35% of the diaphyseal length (from the proximal end, level which was sampled to include fragmentary fossils, see Material and Methods; n=2), *Hapalops* falls in the uppermost distribution of the extant sloths, which does not markedly differ from that of armadillos or anteaters. The CSS at humeral mid-diaphysis distinguishes quite clearly the functional categories, with high values (i.e., elliptical cross-sections) in armadillos, intermediate values in anteaters, and low values (i.e., round cross-sections) in extant sloths. In *Hapalops*, this parameter falls among the particularly tight range of extant sloths (Fig. 3D), but among that of anteaters at 35% of the diaphyseal length. In *Scelidotherium* (n=1), the GC of the humerus at mid-diaphysis is higher than that of most extant xenarthrans, falling in the upper distribution of armadillos and extant sloths (Fig. 3C). One should note, however, that this parameter does not yield any clear distinction among lifestyles. The humeral CSS at mid-diaphysis of *Scelidotherium*, on the other hand, falls among anteaters (Fig. 3D).

There is a clear tendency for the radial diaphysis GC to be higher in armadillos, intermediate in anteaters, and lower in extant sloths. *Hapalops* (n=1; sampled at 72% of diaphyseal length) falls among the distribution of armadillos, being slightly higher than extant sloths’ values (Table 2). The GC of *Glossotherium* and *Lestodon* at radial mid-diaphysis is very low, which agrees with the tendency observed in extant sloths (Fig. 3E). The CSS at that location is found as rather homogenously low among extant xenarthrans, except for two armadillos with outlying high values. *Glossotherium* and *Lestodon* fall beyond the distribution of most extant xenarthrans, their CSS being only tied or exceeded by the two outlying armadillos (Fig. 3F).

Regarding the trabecular architecture parameters, only the degree of anisotropy (DA) will be presented with univariate comparisons, as it was singled out as the most functionally informative of these parameters in extant xenarthrans (Amson et al. 2017a). But mean values of other trabecular parameters of interest are also presented in Table 3. For the humeral head, using a ROI representing 72% of the maximum volume (see Material and Methods section), armadillos are distinguished from other extant xenarthrans by their high values (i.e. more anisotropic architecture). Both *Hapalops* and *Lestodon* (n=1 in each case) fall in the upper distribution (i.e., more anisotropic) of extant sloths and anteaters (Fig. 4A). The same pattern is found for the full ROI in *Lestodon* (no data for *Hapalops*, see Material ad Methods section). In the distal radius (trochlea), the trabecular architecture of armadillos is again found as more anisotropic than in the other extant categories. Moreover, the main distribution of extant sloths is found as clustering at the level of the lower values of anteaters. The DA value of *Hapalops* falls above the main distribution of extant sloths, within that of anteaters (Fig. 4B). *Glossotherium* is the sampled taxon with the single lowest DA value (most isotropic structure). One should note, however, that DA was significantly correlated to body size (see Discussion). The main direction of the trabeculae (MDT) in the radial trochlea (humeral head did not yield lifestyle discrimination; Amson et al. 2017b) of both *Hapalops* and *Glossotherium* falls outside the distribution of extant xenarthrans (Fig. 4C). In both cases, the MDT falls closer to the distribution of extant sloths.

**Figure 4.**
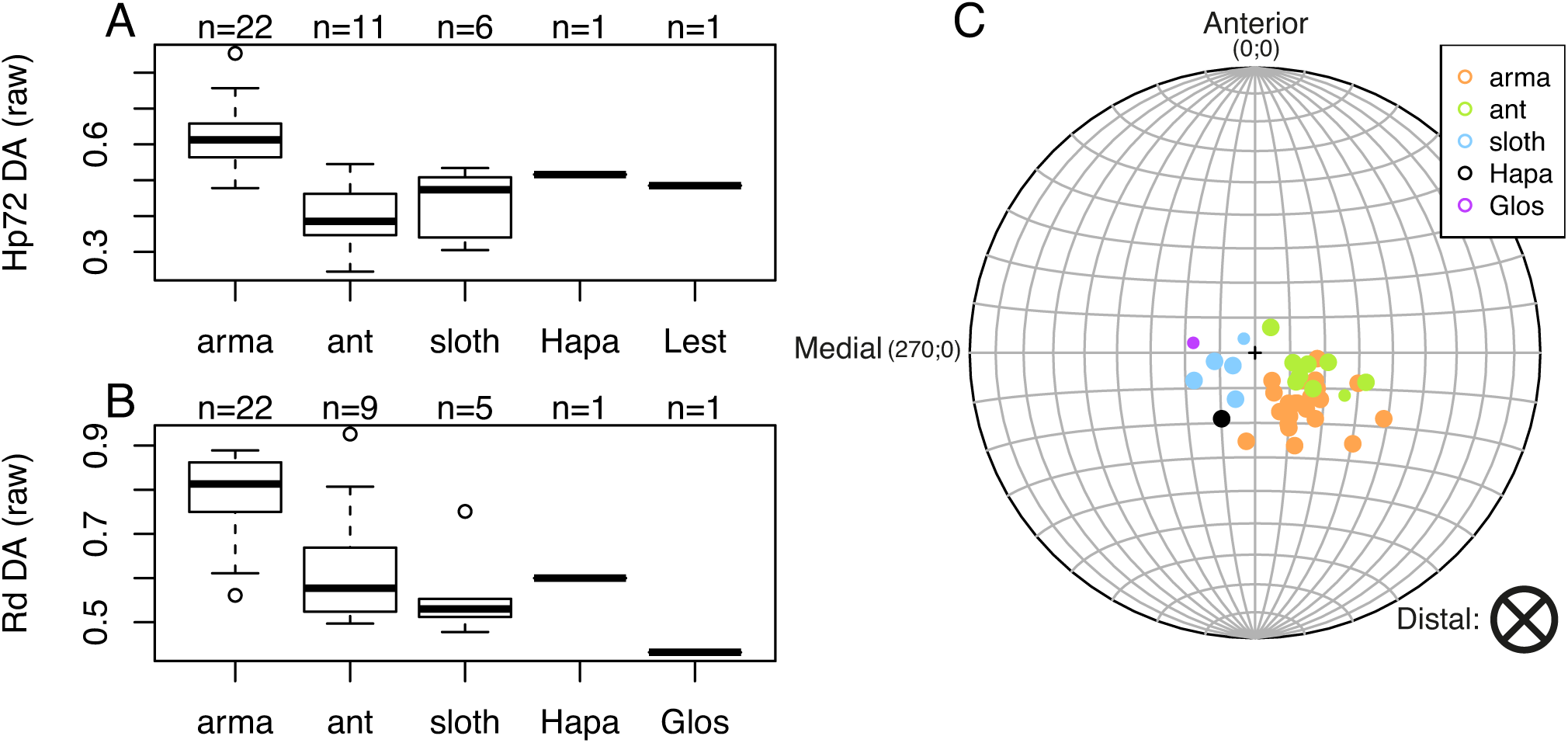
Univariate comparisons of trabecular anisotropy parameters. A, degree of anisotropy (DA) in the humeral head ROI, reduced at 72% of its maximum size (see Material and Methods section); B, DA in the radial trochlea; C, main direction of the trabeculae (MDT) in the radial trochlea. Abbreviations: ant, anteaters; arma, armadillos; Glos, *Glossotherium*; Hapa, *Hapalops*; sloth, extant sloths.

**Table 3.** Mean values of trabecular parameters of interest for each lifestyle category and extinct taxon.

|  | DA (NU) | Conn.D (nb.mm<sup>-3</sup>) | Tb.Th (mm) | Tb.Sp (mm) | BS/TV (mm<sup>-1</sup>) | BV/TV (NU) |
| --- | --- | --- | --- | --- | --- | --- |
| Humeral head 100% |  |  |  |  |  |  |
| Armadillos | 0.60 | 12.35 | 0.25 | 0.47 | 3.38 | 0.41 |
| Anteaters | 0.40 | 11.59 | 0.26 | 0.41 | 3.43 | 0.45 |
| Extant sloths | 0.43 | 9.36 | 0.31 | 0.49 | 3.14 | 0.44 |
| *Lestodon* | 0.50 | 0.58 | 0.80 | 0.81 | 1.23 | 0.58 |
| Humeral head 72% |  |  |  |  |  |  |
| Armadillos | 0.62 | 12.31 | 0.24 | 0.46 | 3.57 | 0.41 |
| Anteaters | 0.40 | 11.38 | 0.26 | 0.42 | 3.56 | 0.45 |
| Extant sloths | 0.44 | 9.03 | 0.32 | 0.50 | 3.16 | 0.45 |
| *Hapalops* | 0.52 | 3.22 | 0.39 | 0.56 | 3.39 | 0.50 |
| Radial trochlea 100% |  |  |  |  |  |  |
| Armadillos | 0.79 | 16.05 | 0.34 | 0.37 | 3.87 | 0.49 |
| Anteaters | 0.63 | 11.19 | 0.30 | 0.44 | 3.20 | 0.44 |
| Extant sloths | 0.56 | 8.75 | 0.30 | 0.52 | 2.39 | 0.40 |
| *Hapalops* | 0.60 | 3.03 | 0.26 | 0.85 | 1.55 | 0.24 |
| *Glossotherium* | 0.43 | 1.02 | 0.43 | 1.25 | 0.59 | 0.23 |

Footnotes. Percentage indicates the cropping coefficient that was used (100% denoting the lack thereof; see Material and Methods section). Abbreviation: NU, no units.

### Phylogenetically flexible discriminant analyses

Each studied “ground sloth” was subject to an independent analysis (see Material and Methods), to predict the most probable lifestyle among the three broad lifestyle categories represented by armadillos, anteaters, and extant sloths, respectively. The results regarding classification of each “ground sloth” are given in Table 4, and the corresponding outcomes of the training data (posterior probability of the classification of the extant species according to each discriminant analysis) are given in SOM 4. We also provide the canonical coefficients (weights) of each explanatory variable for each analysis in SOM 5. For *Hapalops*, 18 parameters could be initially included in the analysis (diaphyseal and trabecular parameters, from both the humerus and radius). Due to high correlation among some variables (Conn.D between two ROIs; between Tb.Th and Tb.Sp of both ROIs; between BS and BV of the radial trochlea ROI), four variables were excluded (see list of included variables in SOM 5. The recovered optimal Lambda is 0 (no significant correlation of the trait values with phylogeny) and the discrimination is optimal (training misclassification error of 0%). *Hapalops* is classified in the category of extant sloths’ lifestyle with a high posterior probability (>99%). Indeed, it falls close to extant sloths’ distribution along the Discriminant Axis (pDA) 1 (Fig. 5A). However, *Hapalops* clearly falls beyond the distribution of extant xenarthrans along pDA2. The parameter contributing the most to the discrimination is the DA (that of the radial trochlea for pDA1 and that of the humeral head for pDA2; see SOM 5).

**Table 4.** Lifestyle classification of the extinct taxa as predicted by phylogenetically flexible discriminant analyses (because of the difference in the included predictive variables for each taxon, a dedicated discriminant analysis was performed for each of them).

|  | Predicted class | P(ant) | P(arma) | P(sloth) |
| --- | --- | --- | --- | --- |
| *Hapalops* | sloth | 0.00 | 0.00 | 1.00 |
| *Lestodon armatus* | arma | 0.35 | 0.64 | 0.01 |
| *Glossotherium robustum* | ant | 0.50 | 0.25 | 0.25 |
| *Scelidotherium leptocephalum* | ant | 0.37 | 0.27 | 0.36 |

Footnotes. Abbreviations: P(“class”), the posterior probability for the extinct taxon to be classified as “class”; ant, anteater; arma, armadillo; sloth, extant sloth.

For *Lestodon*, eight parameters could be included (from the radial diaphysis and humeral head trabeculae), of which one was excluded because of collinearity (present between Tb.Th and Tb.Sp). The recovered optimal Lambda is 0.84, and training misclassification error is 50%. It is classified in the armadillos’ lifestyle category with a rather low posterior probability (64%), the second most probable classification being to anteaters (35%). According to this analysis, a classification in extant sloth’s category is very improbable (0.006%). *Lestodon* falls beyond the distribution of extant xenarthrans (Fig. 5B). The parameter contributing the most to the discrimination is the ‘size-normalized’ Tb.Th (for both pDA1 and pDA2).

**Figure 5.**
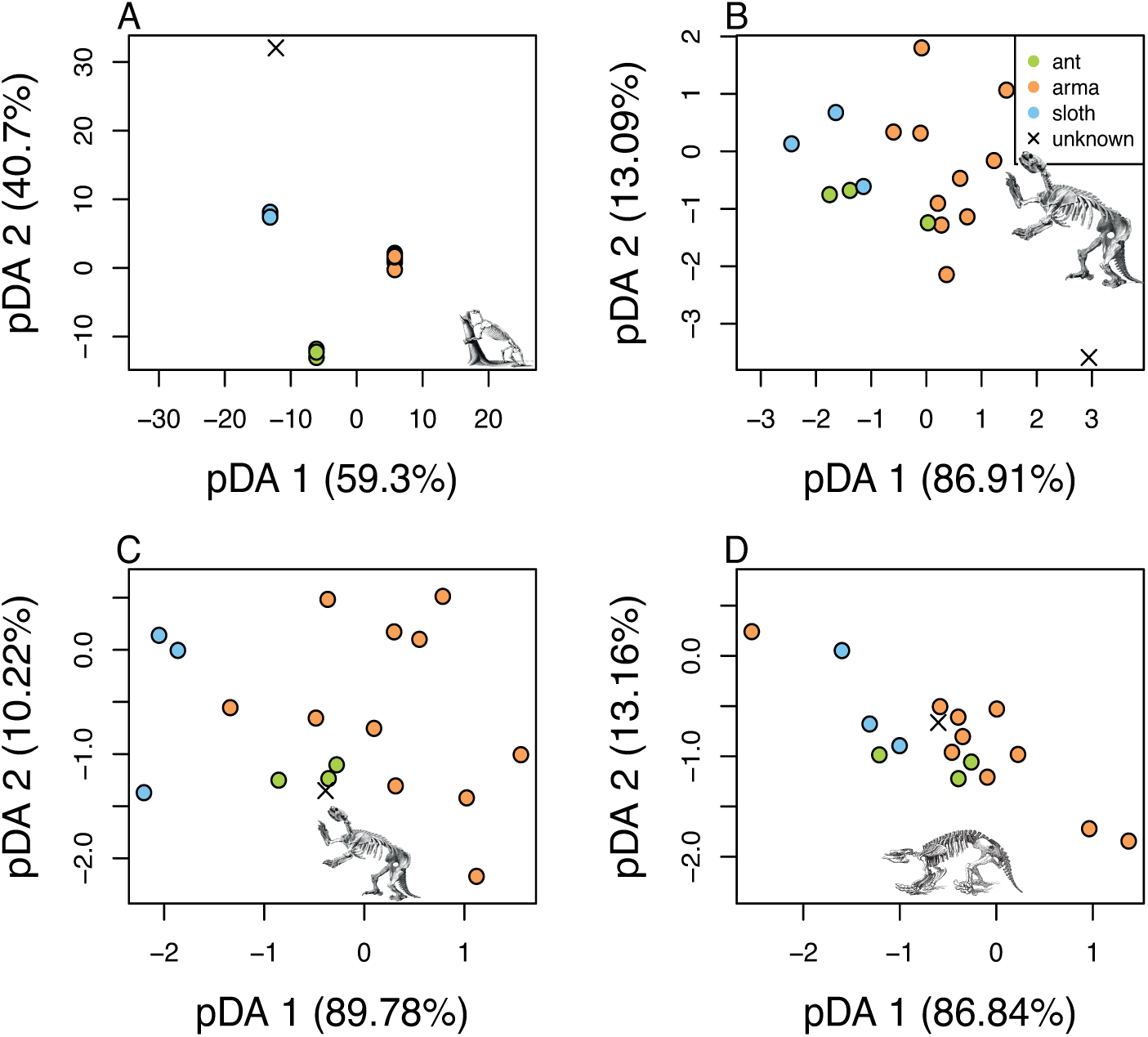
Phylogenetically flexible linear discriminant analyses using humeral and radial bone structure parameters. One analysis per extinct taxon (referred as of “unknown” class) was performed, because of the difference in the parameters that could be included (see Material and Methods section and Table 1). A, *Hapalops*; B, *Lestodon*; C, *Glossotherium*; D, *Scelidotherium*. Abbreviations: ant, anteaters; arma, armadillos; sloth, extant sloths. Next to each discriminant axis is given between brackets the corresponding percentage of explained between-group variance. The size of extinct sloths’ representations gives a rough indication of their body sizes.

For *Glossotherium*, eight parameters could be included (from the radial diaphysis and trabeculae of the radial trochlea). The recovered optimal Lambda is 0.88, and training misclassification error is 35%. The most probable classification is to anteaters (50%), followed by the equally probable classifications to armadillos or extant sloths (each 25%). *Glossotherium* falls within the distribution of extant xenarthrans, but outside the distribution of each lifestyle class, just outside that of anteaters (Fig. 5C). The parameters contributing the most to the discrimination are the DA (pDA1) and ‘size-normalized’ BS (pDA2).

For *Scelidotherium*, only two parameters could be included (from the humeral diaphysis). An optimal Lambda of 0.96 and a high training misclassification error of 69% were recovered. The three possible classifications are roughly equally probable (anteater: 37%; extant sloth: 36%; armadillo: 27%). *Scelidotherium* basically falls in the middle of the distribution of extant xenarthrans (Fig. 5D). The parameter contributing the most to the discrimination is CSS (for both pDA1 and pDA2).

## DISCUSSION

On the whole, the classification of extinct sloths to one of the extant xenarthran lifestyles (that of armadillos, anteaters, or extant sloths) based on forelimb bone structure proved to be challenging. This appears to be due to at least three obvious causes: (1) the imperfect lifestyle discrimination based on diaphyseal and trabecular parameters, (2) the difficulties raised by the size correction (for some parameters), and (3) the fact that the values of extinct taxa are outliers with respect to the distribution of extant xenarthrans (for some parameters).

The four discriminant analyses we performed vary greatly in the number of included parameters. As expected, analyses including more parameters yielded a better discrimination, i.e., a lower misclassification error. The lowest misclassification error (0%) was obtained for the analysis of *Hapalops*, for which it was possible to include 14 parameters (18 before exclusion of collinear parameters) from both the diaphysis and epiphyseal trabeculae. The worst discrimination (69% of misclassification error) was found for the analysis of *Scelidotherium*, for which only two parameters, from the humeral diaphysis, could have been included. This lends support to the approach of combining parameters from several bone compartments, if one endeavours to discriminate lifestyles based on these parameters.

Several of the investigated parameters were significantly correlated with body size. To attempt to prevent the size of the studied taxa from influencing the analysis, a common approach is to size-correct the raw data using the residuals of a regression of the trait against a body size proxy (Mccoy et al. 2006). This proved to be challenging for extinct sloths, because, for most of them, body size largely exceeds that of extant xenarthrans (Vizcaíno et al. 2017). This potentially makes the size regressions spurious, as the extreme values over-influence the regression coefficients. This is not a trivial consideration for our dataset. For instance, if one would size-correct the DA in the radial trochlea using the residuals of the corresponding size regression, the medium-sized extinct sloth *Glossotherium*, of which the raw DA value was found as the lowest of the dataset, would fall in the middle of the overall distribution. For those parameters that are dimensionless, we hence decided to use the untransformed data. But this is likely to be biased as well, due the potential presence of allometry. For instance, the scaling exponent of the degree of anisotropy (DA) across primates in the humeral and femoral head was found by Ryan & Shaw (2013) to be significantly negative (but close to 0, which would have denoted isometry). We also found a negative scaling exponent for one of the investigated ROI, the radial trochlea. It would be suboptimal to exclude this parameter, especially because it was found as the best functionally discriminating parameter in extant xenarthrans (Amson et al. 2017a). It was also singled out as reflecting joint loading in primates better than other parameters (Tsegai et al. 2018), and, more generally, DA was found as functionally informative in several analyses about that clade (e.g. Ryan & Ketcham 2002; Griffin et al. 2010; Barak et al. 2013; Su et al. 2013; Georgiou et al. 2018; Ryan et al. 2018; Tsegai et al. 2018). A tendency for a more anisotropic structure in the femoral head of arboreal squirrels was also demonstrated (Mielke et al. 2018b). A way to improve accuracy of the size-correction using residuals of a regression against a body size proxy would be, in our case, to include to the sampling xenarthrans that have a body size between that of extant species and that of the giant “ground sloths”, i.e., with a mass roughly between 50 kg and 300 kg. Unfortunately, the number of known xenarthrans of this size range is very limited.

It was already obvious from univariate comparisons that the bone structure in *Hapalops*, the small-sized extinct sloth, departed from the condition observed in extant xenarthrans. Indeed, the overall great compactness of its humeral diaphysis does not seem to be matched by any other sampled xenarthran (but see aquatic specialization of *Thalassocnus*; Amson et al. 2014). This does not seem to be a systemic bone mass increase (Amson et al. 2018), because neither the trabecular parameters nor the compactness of the radial diaphysis of this taxon seem to be notably affected by bone mass increase. Finding a compact humerus is particularly surprising, as the stylopod can be expected to be less compact than the zeugopod in terrestrial mammals (Amson & Kolb 2016). In the case of *Lestodon*, it was not obvious from univariate comparisons that its bone structure was outlying, but both the latter and *Hapalops* fell outside the range of extant xenarthrans in the respective discriminant analyses. One may hence conclude that, based on their bone structure, the humerus and radius of both *Hapalops* and *Lestodon* were likely involved in a loading regime different from those associated with the lifestyles of extant xenarthrans. For *Hapalops*, one can however note that the phylogenetically informed discriminant analysis strongly supports a classification within extant sloths’ category, which might indicate that some aspects of their mechanical environment were similar. The main direction of the trabecular (MDT) also agrees with the fact that the bone structure of extant sloths is different from that of *Hapalops*, but that the former represent the most similar of the three extant lifestyles discriminated here (Fig. 4C). Based on bone gross morphology, *Hapalops* was previously reconstructed as partly or primarily arboreal (Matthew 1912; White 1997). Both bone structure and gross morphology therefore seem to point in the same direction for the reconstruction of *Hapalops*’ lifestyle. The large-sized *Lestodon*, on the other hand, is not classified with strong support to one of the extant groups. The least probable classification is to extant sloths’ lifestyle (0.03% of posterior probability), which might suggest that the bone structure of *Lestodon* resembles more that of anteaters and armadillos. Naturally, suspensory posture has never been purported for this elephant-sized sloth. *Lestodon* was interpreted as traviportal (slow-moving with both quadrupedal and bipedal stances) by Toledo (1996), and the forelimb gross morphology was found to be consistent with fossorial activity (but probably not to procure food (Coombs 1983); see Bargo et al. (2000) for a more tempered interpretation). Including other fossorial and non-fossorial taxa in the sampling of the bone structure analysis will be necessary to suggest a more precise assertion regarding the digging habits of this taxon (but its large size might be problematic, see above). The two other extinct sloths subject to a discriminant analysis, *Glossotherium* and *Scelidotherium*, differ from the former two in falling within the distribution of extant xenarthrans. However, in neither case is the classification clear, and it seems that acquiring additional bone structure parameters will be necessary to draw reliable conclusions.

The Mc III did not yield clear discrimination among the extant lifestyles and was hence not included in the discriminant analyses. But one can note that an interesting pattern was observed in the cross-sectional shape (CSS) of extinct sloths at mid-diaphysis. Indeed, high values, denoting elliptic sections, are found in *Valgipes* and *Glossotherium*. Such a bone structure is expected to be suited to resist bending along its major axis (Ruff & Hayes 1983). This is consistent with previous lifestyle reconstruction of *Glossotherium*, which is argued to have had fossorial habits (Coombs 1983; Bargo et al. 2000) supposedly entailing a well-marked main direction of bending. Furthermore, it might suggest that *Valgipes* had similar habits, which, to our knowledge, was never purported.

A medulla filled with spongy bone was observed in large-sized mammals, and argued to be a potential adaptation to graviportality (Houssaye et al. 2015). It does not seem to be possible to easily draw such a conclusion for xenarthrans: whatever their lifestyle, xenarthrans with a mass of roughly 5 kg (e.g., *Tamandua*) and over tend to fill their medullary cavity with spongy bone. This is true for the forelimb, as described here (and as also reported by Houssaye et al. (2015) for the humerus), but likely also for the hind limb: a ‘naturally sectioned’ tibia of the small-sized *Nothrotherium* (less than ca. 100 kg; Amson et al. 2016) reveals that the medullary cavity is entirely filled with dense spongy bone (Fig. 2D). In the case of xenarthrans, the great quantity of diaphyseal trabeculae might be related to another aspect affecting bone structure, such as mineral homeostasis and/or metabolism (Eleazer & Jankauskas 2016; and references therein). While more experimental data is required to discuss it beyond speculation, it was reported that extant sloths (at least the two-toed sloth *Choloepus*) are prone to soft tissue mineralization likely due to mineral imbalance (Han & Garner 2016). One can therefore speculate that the observed great quantity of diaphyseal trabeculae might be a storage mechanism for mineral in excess.

The extremely low metabolism of extant sloths was suggested by Montañez-Rivera et al. (2018) as a potential explaining factor for their low cortical compactness (CC). Indeed, they found that extant sloths depart in that regard from other extant xenarthrans as well as from two extinct sloths (the small-sized *Hapalops* and *Parocnus*). No quantitative assessment of CC was performed here. But we can report that, at mid-diaphysis, the CC of the sampled extinct sloths was generally observed as low (when an observation was possible), similar to armadillos and anteaters. Nevertheless, two specimens showed a rather porous cortex, *Hapalops* (humerus; MNHN.F.SCZ162) and *Glossotherium* (radius; MNHN.F.PAM756), though not as porous as that of most extant sloths. A dedicated analysis of extinct sloths’ CC is required to investigate this trait and possibly use it to inform metabolic rate reconstruction in extinct sloths.

Comparison of long bone’s cross-sections among specimens should be performed at the same location, usually defined as a percentage of the bone’s length (e.g., Ruff & Hayes 1983). Here, mid-diaphysis (i.e., 50% of bone length) was selected for complete bones, and, for fragmentary specimens (some fossils), it is the preserved level closest to mid-diaphysis that was used (the other specimens were resampled accordingly). Because of the xenarthran bones’ morphology, most examined cross-sections were located at the level of a prominent bony process. One could therefore consider selecting cross-sections avoiding those processes to test their influence on bone structural parameters. Acquiring cross-sectional properties along the whole diaphysis and assessing the proximodistal evolution of biomechanical properties can also be considered for complete bones (Houssaye & Botton-Divet 2018).

In addition to lifestyle, one can expect that the factors affecting bone structure are the individual’s age, health status, and possibly other features varying intraspecifically (such as sex differences; Eckstein et al. 2007). Details regarding these potential factors are mostly unknown for fossils (and often for recent specimens as well). To control for these factors as much as feasible, the sampled specimens were chosen to be devoid of apparent bone diseases and skeletally mature (even though several presented a remnant of epiphyseal line, see above). It is our assumption that variations in bone structure that relate to a different lifestyle can be expected to be of greater magnitude than intraspecific variations. But this chiefly remains to be demonstrated.

## CONCLUSION

Bone structure of the diaphysis and epiphyses of the third metacarpal, humerus, and radius was here investigated in several species of extinct sloths, comparing it to that of extant xenarthrans. Related parameters were successfully acquired and included in phylogenetically flexible discriminant analyses. The latter constitute, to our knowledge, the first analyses that conjointly include both diaphyseal and trabeculae parameters to discriminate lifestyles. However, no extinct sloths are here confidently ascribed to one of the lifestyles exhibited by extant xenarthrans. This might be due to several factors, and we identified as challenges for the present analysis the lack of discrimination power of some parameters, the difficulties raised by size-correlated parameters, and the fact that some parameters fall outside the range described by extant taxa. The humeral and radial structure of the small-sized *Hapalops*, from the Miocene of Argentina, was nevertheless found as more reminiscent of that of extant sloths, which agrees with the conclusions drawn based on gross morphology. The humeral and radial structure of the large-sized *Lestodon*, from the Pleistocene of Argentina, clearly departs from that of extant sloths, and is more similar to that of anteaters and armadillos. The singular bone structure of xenarthrans, including a medullary cavity filled with spongy bone in most taxa, and a low cortical compactness in extant sloths, deserves further investigation. Because Xenarthra is argued to be one of the four early diverging clades of placental mammals (Delsuc & Douzery 2008; Asher et al. 2009; Gaudin & Croft 2015), such investigations are not only important for the understanding of the evolutionary history of the clade, but potentially for that of Mammalia as well.

## AKNOWLEDGEMENTS

We warmly thank the following curators and collection managers: Guillaume Billet (Muséum national d’Histoire naturelle, Paris; MNHN), Cástor Cartelle (Museu de Ciencias Naturais da Pontifícia Universidade Católica de Minas Gerais, Belo Horizonte, Brazil), Anneke H. van Heteren (Zoologische Staatssammlung München), Frieder Mayer and Christiane Funk (Museum für Naturkunde Berlin), Thomas Kaiser and Nelson Ribeiro Mascarenhas (Universität Hamburg), Irina Ruf and Katrin Krohmann (Senckenberg Forschungsinstitut und Naturmuseum), Stefan Merker (Staatlichen Museums für Naturkunde Stuttgart), Eva Bärmann (Zoologische Forschungsmuseum Alexander Koenig). Anneke H. van Heteren (Zoologische Staatssammlung München), Patrick Arnold (Friedrich-Schiller-Universität Jena), and Aurore Canoville (NC State University) are acknowledged for helping with acquisition of the extant species scans. We thank Patricia Wills, Marta Bellato, and Maïté Adam (AST-RX platform, MNHN) for acquiring the extinct species scans. We acknowledge Luis D. Verde Arregoitia for his help with the function pFDA. Andrew Pitsillides (acting as a reviewer), an anonymous reviewer, Alexandra Houssaye (acting as a PCI Paleo Recommender) and an additional PCI Paleo Recommender are thanked for the improvement they brought to the manuscript.

## Institutional abbreviations

MCL: Museu de Ciencias Naturais da Pontifícia Universidade Católica de Minas Gerais, Belo Horizonte, Brazil;
MNHN.F: Muséum national d’Histoire naturelle, Paris, France, Palaeontology collection;
ZMB_MAM: Museum für Naturkunde Berlin (Germany), Mammals Collection;
ZSM: Zoologische Staatssammlung München, Germany.

## ADDITIONAL INFORMATION

### Author contributions

Conceptualization and methodology, EA, JAN; Formal analysis, EA; Investigation, EA; Writing – Original draft, EA; Writing – Review & editing, EA, JAN. All authors gave final approval for publication.

### Data availability

All the raw scans of fossil specimens sampled for the present analysis will be available from the MNHN collection database pending an embargo. The extant species specimens sampled come from various collections. The corresponding raw scans are available upon reasonable request to the authors.

### Supplementary information

SOM 1. Raw data. Excel document, of which each worksheet corresponds to a sampled region.

SOM 2. ImageJ macro to crop isometrically a stack in 3D.

SOM 3. ImageJ macro to acquire mid-diaphyseal parameters.

SOM 4. Lifestyle classification of the extant taxa, the training data of the phylogenetically flexible discriminant analyses (one analysis was performed per extinct taxon, each on a different worksheet).

SOM 5. Canonical coefficients for each phylogenetically flexible discriminant analysis (one analysis was performed per extinct taxon, each on a different worksheet).

